# TRAP-based allelic translation efficiency imbalance analysis to identify genetic regulation of ribosome occupancy in specific cell types *in vivo*

**DOI:** 10.1101/2020.08.24.265389

**Authors:** Yating Liu, Anthony D. Fischer, Celine L. St. Pierre, Juan F. Macias-Velasco, Heather A. Lawson, Joseph D. Dougherty

## Abstract

The alteration of gene expression due to variations in the sequences of transcriptional regulatory elements has been a focus of substantial inquiry in humans and model organisms. However, less is known about the extent to which natural variation contributes to post-transcriptional regulation. Allelic Expression Imbalance (AEI) is a classical approach for studying the association of specific haplotypes with relative changes in transcript abundance. Here, we piloted a new TRAP based approach to associate genetic variation with transcript occupancy on ribosomes in specific cell types, to determine if it will allow examination of Allelic Translation Imbalance (ATI), and Allelic Translation Efficiency Imbalance, using as a test case mouse astrocytes *in vivo*. We show that most changes of the mRNA levels on ribosomes were reflected in transcript abundance, though ∼1.5% of transcripts have variants that clearly alter loading onto ribosomes orthogonally to transcript levels. These variants were often in conserved residues and altered sequences known to regulate translation such as upstream ORFs, PolyA sites, and predicted miRNA binding sites. Such variants were also common in transcripts showing altered abundance, suggesting some genetic regulation of gene expression may function through post-transcriptional mechanisms. Overall, our work shows that naturally occurring genetic variants can impact ribosome occupancy in astrocytes *in vivo* and suggests that mechanisms may also play a role in genetic contributions to disease.

## Introduction

Proper regulation of gene expression is essential to normal biological function. Sequence variations in key regulatory elements such as promoters and enhancers can alter DNA-protein interactions, and thus alter the transcription of target genes. Such genomic variants have been shown to be statistically over-represented in disease associations (Carullo and Day, 2019; Karnuta and Scacheri, 2018; Schoenfelder and Fraser, 2019). However, final expression of genes, especially protein coding genes, is a multistep process with post-transcriptional regulation also substantially modifying the amount of protein produced during translation of specific transcripts by the ribosome. As with DNA sequence variation, alterations in the sequence of the mRNA can similarly alter RNA-protein or RNA-RNA interactions, thus substantially modulating levels of translation. Key examples include alterations of the Kozak consensus sequence around start codons, the presence of upstream open reading frames, and other sequences in 5’ untranslated regions (UTRs) that can modulate translation initiation (Leppek et al., 2018), as well as polyadenylation signals and miRNA and RNA-binding protein (RBPs) sites in 3’UTR that can influence transcript stability, localization, and translation (Mayr, 2017). However, little attention has been paid to how naturally occurring genomic sequence variation can contribute to modulating ribosome occupancy of specific transcripts (Hou et al., 2015). Additionally, mRNA abundance and stability are not always clear indicators of ultimate protein synthesis; relative mRNA expression/stability and translational levels for a given gene can vary significantly, owing to a host of post-transcriptional events (Liu et al., 2016). Indeed, assessment of Translational Efficiency (TE), defined empirically as the relative number of ribosomes per transcript using combinations of Ribosome Footprinting and RNA sequencing (Dalal et al., 2017; Ingolia et al., 2009, 2011) has suggested that transcript abundance only accounts for 43%-63% of ribosome occupancy across transcripts. Further, post-transcriptional regulation is especially important in the CNS, where both neurons and astrocytes show evidence of complex activity dependent and localized regulation of translation (Sakers et al., 2017; Sapkota et al., 2020). To what extent do variants in UTRs also have substantial impact on final levels of ribosomes on the mRNA? Do they also have the propensity to contribute to normal variation across a population? Does this occur *in vivo* and in tissues, such as the brain, where translation regulation is highly sophisticated?

To address these and similar questions, we aimed to develop an approach for assessing the influence of genetic variation on ribosome occupancy in specific cells in the brain. We adapted a classical approach to the association of genetic variation to transcript abundance: Allelic Expression Imbalance (AEI). Allelic Expression Imbalance is a very sensitive within-subject design for associating a particular genetic haplotype with an imbalance of in transcript abundance, such as might be produced by alteration in transcription due to variants in enhancers nearby a gene. It works by analyzing the relative abundance of tag-SNPs present in an mRNA from heterozygous alleles. It has been widely used in humans (Chuang et al., 2017; Mohammadi et al., 2019) and model organisms (Chen et al., 2019; Pinter et al., 2015; Zhuo et al., 2017) to study genetic regulation of transcription, as well as more unusual phenomena such as imprinting (Chuang et al., 2017; Santoni et al., 2017), parent of origin effects, and allele-specific protein-RNA binding differences (Bahrami-Samani and Xing, 2019). To study genetic regulation of translation *in vivo*, we combined this framework with Translating Ribosome Affinity Purification (Dougherty, 2017; Heiman et al., 2008) - a method for purifying the mRNA specifically bound to ribosomes from genetically targeted cell populations. TRAP has been shown to reflect dynamic changes in translation (e.g. by iron response elements) in a way that quantitatively matched polysome fractionation (Heiman et al., 2008). By collecting in parallel total mRNA-seq as well as ribosome bound mRNA-seq from the same hybrid mice, we reasoned we could simultaneously analyze AEI, Allelic Translation Imbalance (ATI), and potentially Allelic Translational Efficiency Imbalance (ATEI) to identify those alleles where ribosome occupancy was significantly altered from what was expected from initial transcript abundance. This builds upon a similar study using polysome profiling and ribosome footprinting in cultured cells to examine ATI and AETI (Hou et al., 2015). The TRAP method could be advantageous because it simplifies purifying actively translating mRNAs from cell lysate as compared to more lengthy approaches such as polysomal profiling or ribosome footprinting. More importantly, the method also allows uniquely cell-specific capture of the translating pool from *in vivo* specimens. Because we are interested in astrocytes, it is employed here using an astrocyte-specific promoter-driven expression of the TRAP construct as a test case. A Cre-inducible TRAP mouse could be mated to virtually any cell-type specific Cre expression mouse for targeted allele-specific translatomics. Further, since ribosomes are assembled in the nucleolus resulting in a speck of nuclear GFP, the same construct can be used to collect nuclear RNA from the same cell type by parallel in FACs (Reddy et al., 2017). Though not done here in this first test of the method, having a more cell type specific AEI to compare to the ATI could further enhance the sensitivity of the approach.

The prior work in cultured cells has shown that F1 hybrids of *Mus musculus* and the related *Mus spretus* have ∼1000 genes which show ATEI (roughly 14% of measurable transcripts), and that SNPs in the 5’ UTR, especially those close to the start codon, contributed significantly to this, as well as SNPs in this region that caused predicted less stable secondary structure (Hou et al., 2015). A similarly constructed follow-up study demonstrated ∼600 genes with allelic decay imbalance, with increased instability owing largely to a greater presence of miRNA sites, exclusively in the 3’UTR (Sun et al., 2018).

Here, we show that our similar *in vivo* approach can identify thousands of variants that are specifically associated with alterations in ribosome occupancy. We found thousands of haplotypes associated with AEI, and a slightly larger number that altered ATI, as well as 138 that specifically modulated ribosome occupancy through ATEI. Finally, by annotating all variants based on their alteration of putative translation regulatory sequences in UTRs, we highlight numerous examples of ATEI genes with variants altering uORFs, polyA signals, and notably the strength of microRNA binding sites.

## Methods

### Mouse Husbandry

All animal procedures were approved by the Washington University Institutional Animal Care and Use Committee. Aldh1L1 TRAP mice (Tg(Aldh1l1-EGFP/Rpl10a)JD130Htz, JAX:030247) originally generated as BAC transgenics on an FVB background, were backcrossed >12 generations to C57BL/6J mice (hereafter BL6). Males of this line were then crossed to female FVB mice to generate N1 hybrids. Two male and two female N1 littermates (N = 4) were genotyped for the presence of the TRAP allele and sacrificed at P21 for TRAP-Seq.

### Immunofluorescence

Following euthanasia, Ald1h1L1 TRAP mice were perfused with phosphate buffered saline (PBS), and then 4% paraformaldehyde in PBS. Brains were cryoprotected in PBS sucrose solution, then frozen in OCT and then cryosectioned into 40 micron floating sections and stored in PBS 0.25% sodium azide at 4C until use. For assessment of astrocyte labeling, sections were blocked and solubilized with 5% normal serum and 0.25% triton X-100 in PBS, then incubated with Chicken anti-GFP (ab13970, abcam), and Goat anti-Glt1 (sc-7760, Santa Cruz biotechnology), and stained with appropriate Alexa fluor conjugated secondary antibodies, and counterstained with DAPI. Slices were slide mounted in Prolong Antifade, and imaged by confocal microscopy.

### TRAP and RNASeq

TRAP was performed on P21 whole brain homogenates as described previously (Heiman et al. 2008 and Sapkota et al. 2020). RNA was extracted from the input and TRAP samples using the RNA Clean & Concentrator Kit (Zymo Research). RNA quality and concentration were measured using a NanoDrop spectrophotometer and a High Sensitivity RNA ScreenTape (Agilent Technologies). All samples had RNA integrity scores >7.0.

Barcoded libraries were prepared with the NEBNext Ultra II RNA Library Prep Kit for Illumina (NEB #E7770) and the NEBNext rRNA Depletion Kit (NEB #E6310), according to the manufacturers’ instructions. Each library was screened for quality and adapter dimers on a High Sensitivity D1000 ScreenTape (Agilent Technologies), then sequenced on an Illumina HiSeq 3000 using 150bp, single-end reads.

Data is available at GSE156414.

### Standard TRAP and RNASeq analysis

Sequencing results were quality checked using FastQC (version 0.11.7). Illumina sequencing adaptors were removed using Trimmomatic (version 0.38) (Bolger et al., 2014), and reads aligning to the mouse rRNAs were removed using bowtie2 (version 2.3.5) (Langmead and Salzberg, 2012). Surviving reads were then aligned to the mouse transcriptome (Ensembl Release 93) using STAR (version 2.7.0d) (Dobin et al., 2013). The number of reads mapped to each gene were counted using htseq-count (version 0.9.1).

Differential expression analysis was done using edgeR (version 3.24.3) (Robinson et al., 2010). Only genes with > 1 CPM in at least 4 out of 8 samples were retained for further analysis (13,645 genes). A negative binomial generalized log-linear model (GLM) was fit to the counts for each gene. Likelihood ratio tests (LRT) were conducted for comparing RNASeq samples with TRAPSeq samples. Known astrocyte genes are defined as described in (Dougherty et al., 2012).

### Genome and annotation

FVB genome from NCBI (https://www.ncbi.nlm.nih.gov/assembly/GCA_001624535.1) and BL6 genome from Ensembl (GRCm38.p4) were merged to construct a pseudo-genome for allele specific alignment. Sex chromosomes X and Y were removed from the final pseudo-genome. Pseudo-annotation was also constructed by combining FVB annotation from UCSC mouse strain assembly hub (Ensembl version 86) and BL6 annotation (Ensembl version 86). Only genes existing in both FVB and BL6 were kept in the final pseudo-annotation.

### Allele-specific expression analysis

Reads that passed the quality check and did not align to the mouse rRNAs, as above, were aligned to the pseudo-genome with multimapping disallowed using STAR (version 2.7.0d) (Dobin et al., 2013). This resulted in only reads containing SNPs, thus enabling unique alignment to either FVB or BL6 sequences, being retained. The exon regions were extracted from the pseudo-annotation. The number of reads overlapped with each exon were counted using bedtools (version 2.27.1). Gene counts were calculated by summing up the counts of their exons. From these counts, each sample was divided into two pseudo-samples representing its FVB and BL6 counts for each gene for the purposes of edgeR analysis.

### Library complexity

Gene counts lower than 20 were removed and remaining counts were normalized based on upper quartile normalization. The BL6 expression bias of each gene was calculated as the proportion of BL6 normalized counts from the total allele specific normalized counts. Library complexity was measured by fitting a beta-binomial distribution of BL6 bias using the VGAM package. The shape parameters, a and b, of beta-binomial distributions were estimated. The dispersion values (r < 0.05) were used to indicate whether the libraries are sufficiently complex. The library complexities were ranged from 0.056 – 0.071 (Fig. S2).

### AEI, ATI, and ATEI analyses

All raw gene counts were input to edgeR (version 3.28.0) for differential expression analysis. Only genes with > 1 CPM in at least 4 out of 8 samples were retained for further analysis (9,021 genes). A negative binomial generalized log-linear model (GLM) was fit to the counts for each gene. Then the likelihood ratio tests (LRT) were conducted for comparing allelic expression, allelic translation, and allelic translation efficiency. The comparison models are listed as below:

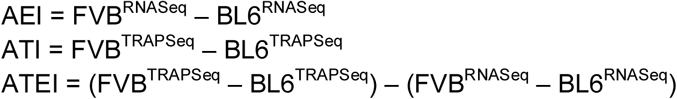

Differentially expressed genes in each category were defined by having FDR < 0.1. Results of all comparisons are included in Supplemental Tables 2 and 3.

### UTR annotation pipeline

There are a few feature elements in non-coding regions that are known as translation regulators, including the number of AUGs in the 5’ UTR, the number of Kozak sequences in the 5’ UTR, the number of polyA signals in the 3’ UTR, the stop and start codon, and the Kozak score at the transcription initiation site (TIS). In order to identify the potential deleterious variants which could disrupt those translation regulators in non-coding sequences, a UTR annotation pipeline (Fig. 4A) was developed to annotate variants on whether and how it alters any of these feature elements in UTRs. As non-coding sequences playing a role in gene regulation may be conserved within a genome, the UTR annotation pipeline also queried conservation scores of variants’ positions to identify variants overlapping with highly conserved regions in the downstream analysis. The conservation scores are from the UCSC phastCons60way and phyloP60way. The Position Weight Matrix to determine the Kozak score was derived from data in an empirical study (Sample et al., 2019), graciously provided by the authors.

### Annotation of Variants

FVB specific variants relative to the reference mouse genome BL6 were obtained from the Mouse Genomes Project SNP and indel Release Version 5 (ftp://ftp-mouse.sanger.ac.uk/REL-1505-SNPs_Indels/) (Keane et al., 2011). Variants were annotated using the UTR annotation pipeline to identify variants that disrupt the feature elements in non-coding regions. Gene models from Ensembl release 80 were used for annotation.

### MiRNA analysis

Variants that overlapped with the ATEI genes were selected for miRNA analysis. For each variant, 34 nucleotides up- and downstream of the varying sequence was pulled from the BL6 genome as BL6 mRNA. The corresponding FVB mRNA was generated using BL6 mRNA and variant information. BL6 mRNA and FVB mRNA were separately input into miRanda v3.3a (Enright et al., 2003) to predict their target miRNAs with score threshold as 80 and energy threshold as −14 kcal/mol. In order to compare the miRNA target difference between BL6 and FVB caused by the variant, miRNAs that bind to regions outside of variant positions were filtered out. Then the minimum free energy (MFE) of each miRNA from BL6 and FVB were compared, and miRNAs were called only if their absolute MFE difference is larger than 0 (Fig. 4C).

### Code Availability

Code is available on bitbucket: https://bitbucket.org/jdlabteam/allelic_translation_imbalance/src/master/.

## Results

### Development of an *in vivo* approach to identify transcripts with allelic translational efficiency imbalances

Our goal was to measure ribosome bound transcripts from a specific cell type in the brain, and to determine the magnitude of impact of common genetic variation on transcript ribosome occupancy. To achieve this, we crossed two fully sequenced common strains of inbred mice (Fig. 1A), differing with 6,617,019 known small nucleotide variants (SNVs), including SNPs and indels, 110,559 of which occur in mature transcripts (Fig. 1B). Overall, 2354 genes only differ by one SNV in the transcript. We selected these lines rather than the more distantly related *Mus spretus* to 1) have less variants per transcript, making it more likely the remaining variants were causal, and 2) to model the magnitude of variation that might be seen between two individuals of the same species, such as across individual humans. We included the astrocyte TRAP allele, which enabled purification of ribosomes specifically from astrocytes in the brain (Fig. 1C), and conducted both RNASeq and TRAPSeq from the same lysates. Immunofluorescence analysis revealed that GFP-RPL10A expression was mainly localized to astrocytes as seen by colocalization with the astrocyte marker GLT1 (Fig. 1D). Quality control analyses confirmed that TRAP was reproducible across replicates (Fig. 1E), and that it enriched specifically for known astrocyte transcripts as expected (Fig. 1F).

**Figure 1:**
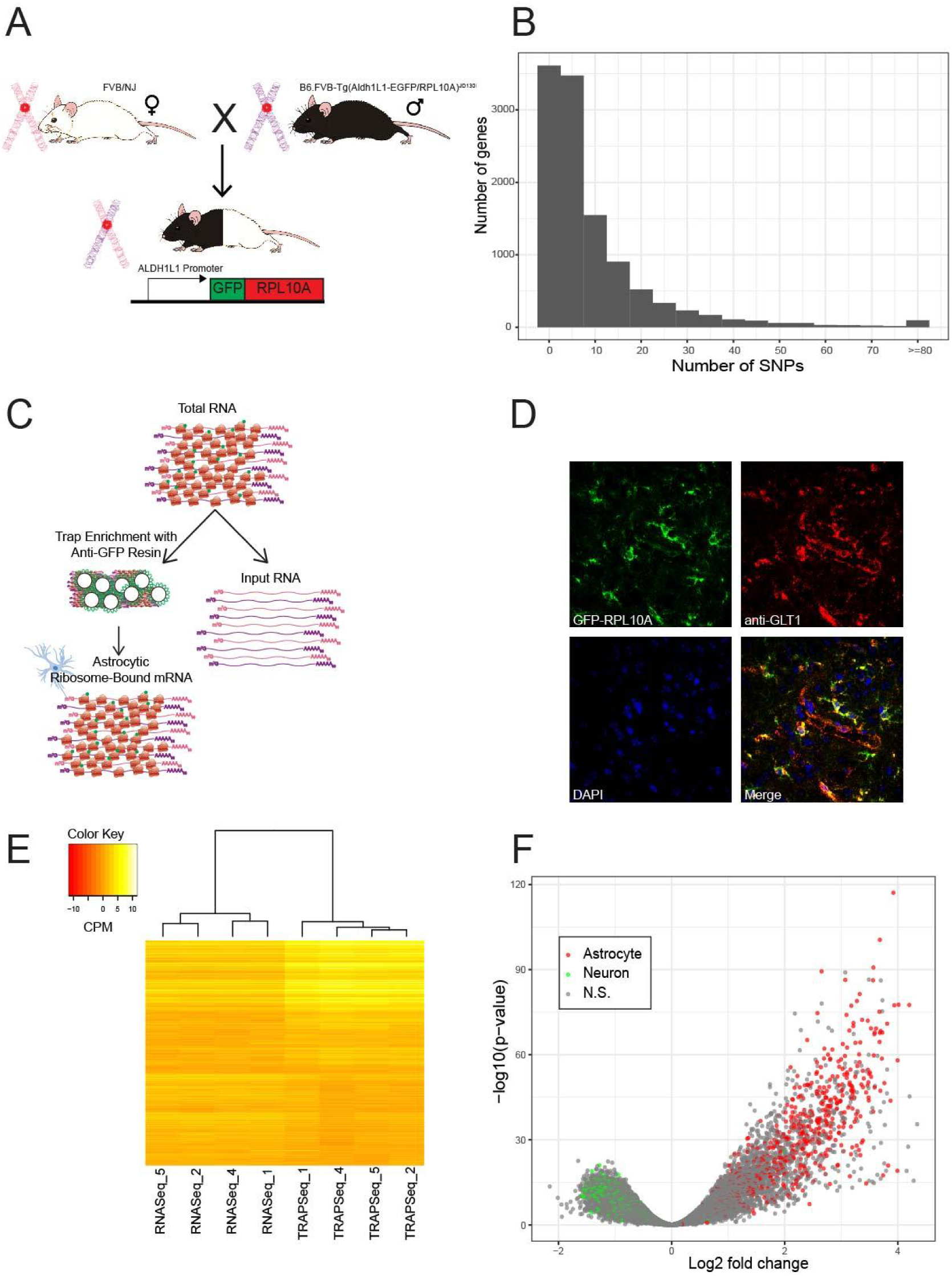
Experimental approach and confirmation of astrocyte TRAP. A) Schematic of experimental design: BL6 with the Aldh1L1 TRAP allele and FVB mice were crossed to generate F1 hybrids with one copy of each strain’s allele. B) Histogram of numbers of transcript-encoded variants per mature gene transcript, between BL6 and FVB. C) Schematic of TRAP - cell type specific expression of a GFP tagged ribosomal protein in astrocytes enables enrichment of ribosome-bound transcripts from these cells. D) Immunofluorescence of cortical sections showing GFP-RPL10A, astrocyte-specific marker GLT1 (red), DAPI, and merged panels showing colocalization, indicating the TRAP construct is expressed in GLT1 positive astrocytes. E) A heatmap and hierarchical clustering across all genes shows that TRAPSeq and RNASeq clustered separately from each other, but were reproducible within groups. F) TRAPSeq showed the expected enrichment of known astrocyte marker genes (red) when compared to standard RNASeq.

We then separately counted transcript fragments that mapped unambiguously to either the FVB or BL6 from each sample, using an efficient new approach that adapts standard RNAseq tools. Specifically, we structured the data as strain-specific ‘pseudo-samples’ in a manner that allowed us to readily adapt edgeR (3.28.0) RNASeq package assumptions to measures of AEI (from RNASeq), as well as ATI (from TRAPSeq) (Fig. 2, Table S1: raw counts, by sample by parent). This further enabled us to sensitively identify ATEI, again using an edgeR framework, as transcripts where the TRAPSeq allelic differences were not proportional to RNASeq allelic differences, suggesting alteration of translation efficiency *in vivo* (Fig 2F). Quality control analyses confirmed that pseudo-samples clustered well by both TRAP and strain (Fig. 3A), showing strong reproducibility. Our approach performed reasonably well, with both AEI and ATI showing roughly equal proportions of variant reads from each BL6/FVB genome (Fig. S2). Thus, we were able to identify 3505 and 3762 transcripts showing AEI and ATI respectively (FDR < .1, Fig. 3B,C, Tables S2 and S3). Correlation between ATI and AEI overall was strong (r = 0.95, Fig. 3D), indicating that the majority of AEI from differences in transcription/transcript abundance carries over to change ribosome occupancy. However, we did detect 138 transcripts where ATEI clearly occurs (FDR < 0.1, Fig 3D,E, and Fig. S1 for top 15 ATEI genes). Overall it appears common variants impact translation efficiency in 1.53% of transcripts in astrocytes under these conditions. This suggests a proportion of transcripts show such translational regulation, and that this type of variation could contribute to final protein expression levels (and occasionally serve as a mechanism for disease associations). However, it is worth noting TRAPSeq in the astrocyte and RNASeq in the whole brain comparisons may not be entirely equivalent, and some very specific changes in astrocytes might be masked. Regardless, Gene Ontologies analysis using DAVID (Huang et al., 2009) failed to highlight any significant categories after correction, suggesting this class of regulation is not specific to any particular biological processes.

**Figure 2:**
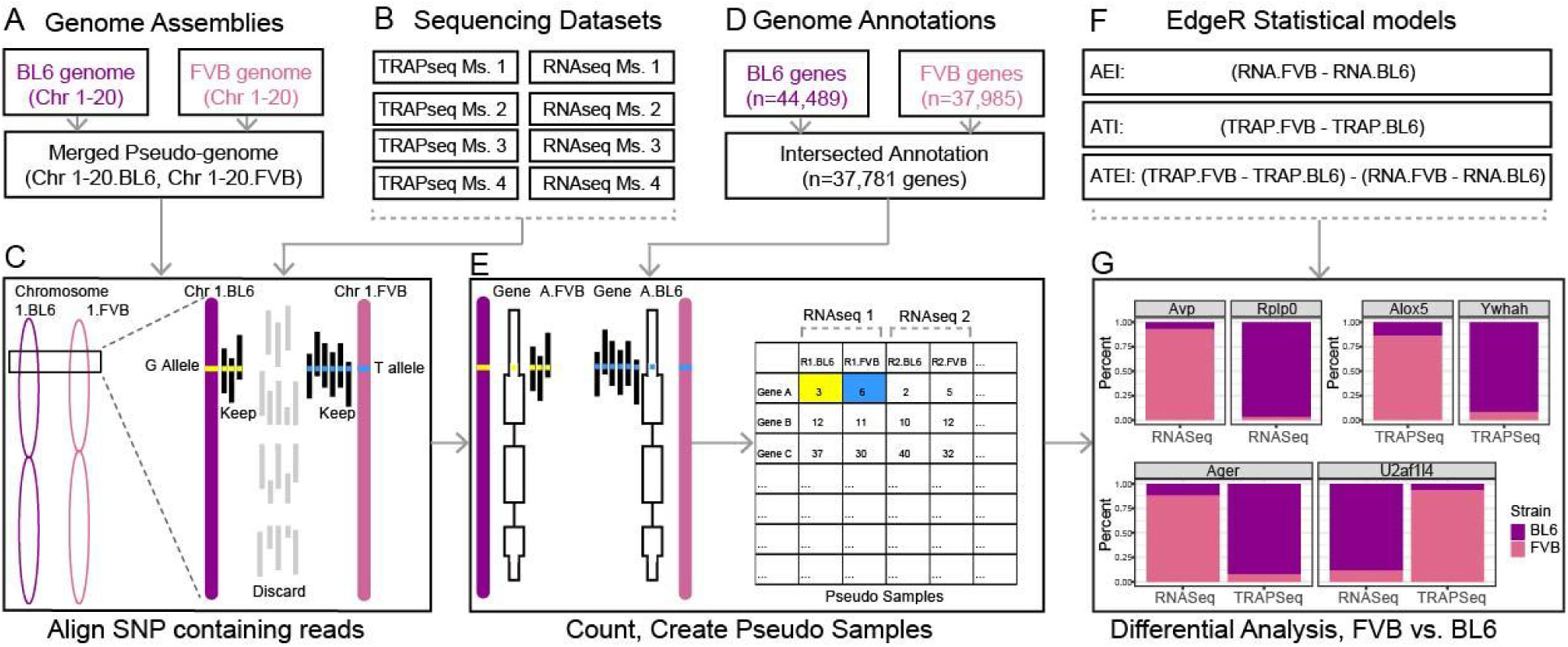
Schematic for analytical approach to identify AEI, ATI, and ATEI. A) BL6 and FVB genome sequences were concatenated into a single genome file (i.e. with 40 chromosomes). B) TRAPSeq and RNASeq reads were aligned to this pseudo genome. C) Only unique alignments were kept, thus discarding any reads lacking SNPs that would distinguish FVB from B6 alleles. D) A merged annotation was also assembled from the two strains, and following counting reads aligning to each gene, E) each sample was divided into two ‘pseudo-samples’ by separating the reads aligning to the B6 and FVB isoform of each gene. F) Statistical models used for EdgeR to compare pseudosamples to detect AEI, ATI, and ATEI, resulting in G) detection of genes significantly different between strains in their expression or ribosome occupancy. These are illustrated with stacked bars showing the percent of reads coming from either the BL6 (purple) or FVB (pink) genomes. Top row shows examples of AEI, ATI transcripts, and bottom row examples of ATEI, where TRAPSeq is significantly diverged from RNASeq.

**Figure 3:**
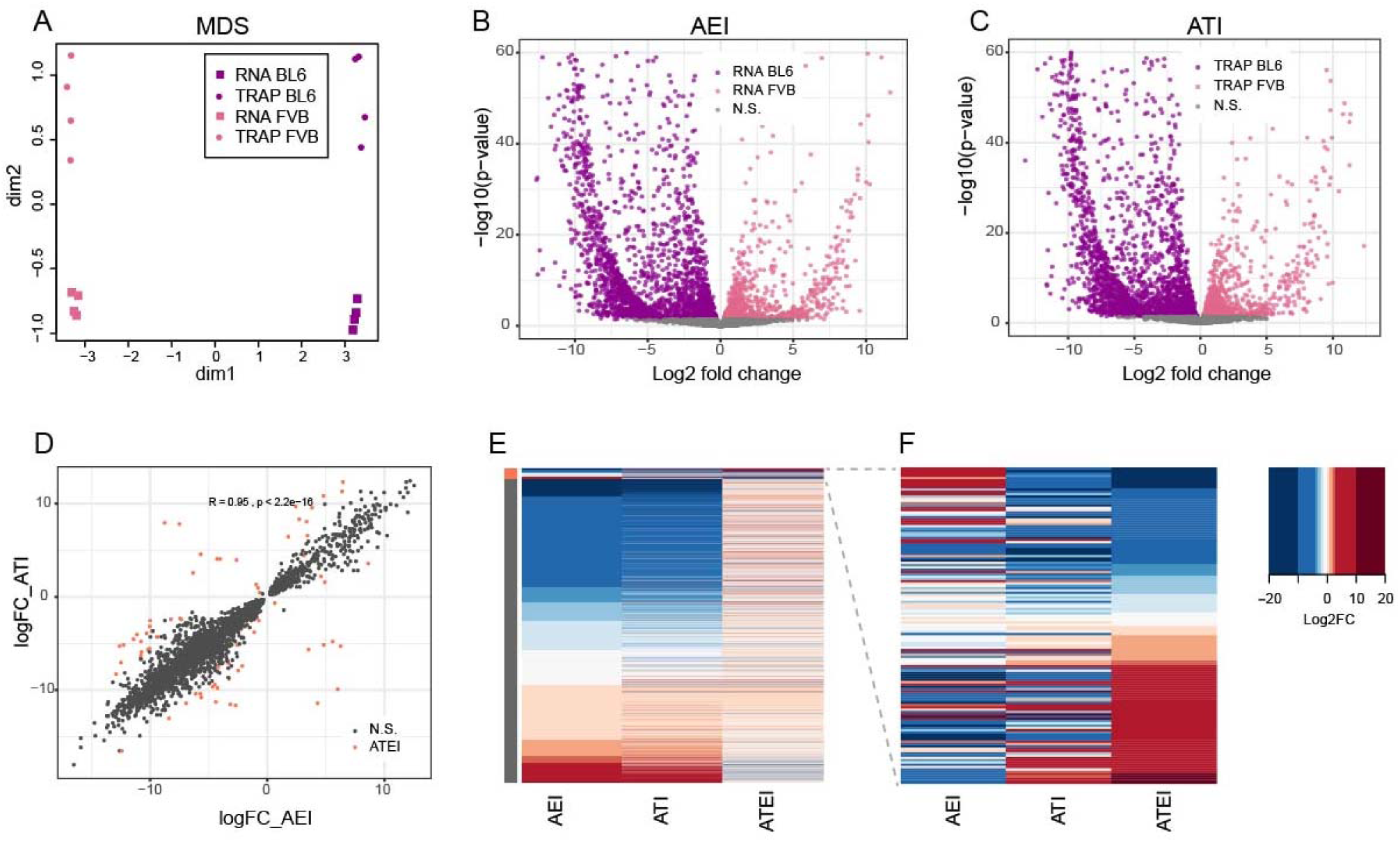
Genetic variation drives substantial changes in RNA abundance and translation. A) Multidimensional scaling plot of pseudo-samples shows clear separation between TRAP and Pre-TRAP horizontally, as well as FVB and B6 allele expression vertically, highlighting reproducibility of replicates and robust differences between strains and sample types. B) Volcano plot for AEI transcripts shows substantial RNA abundance imbalances between strains (*pink: genes where FVB allele is expressed higher, purple: genes where FVB allele is expressed significantly higher*) C) Volcano plot for ATI identified transcripts showing a similar scale of allelic imbalances on astrocyte ribosomes by TRAP. D) Comparisons of ATI and AEI fold changes between B6 and FVB alleles shows that for most genes RNA abundance predicts T AP, though individual genes (orange) show significant ATEI. E) Heatmap of individual transcripts showing strain specific differences in AEI, ATI, and ATEI translation efficiency, including the union of AEI, ATI, and ATEI significant transcripts. The left horizontal side bar indicates ATEI significant transcripts (orange) and ATEI non-significant transcripts (grey). F) Inset highlighting the ATEI significant transcripts.

### Variants that alter uORFs and poly-Adenylation signals occur more often in ATEI transcripts

We next developed a pipeline to identify potential molecular consequences of variations in UTRs (Fig. 4A), enabling us to test several hypotheses regarding which classes of variants might be enriched in ATEI-exhibiting transcripts. A variety of sequences such as upstream open reading frames (uORFs), Kozak consensus sequences (Kozaks), Start and Stop Codons, as well as PolyAdenylation signals (PAs) could all plausibly alter TE. We scanned all variant-containing transcripts to determine whether any common variants altered any of these, and if this occurred more often in ATEI transcripts than expected by chance. We found that uORFs and polyA signals were altered more often in ATEI transcripts than expected by chance (odds ratio > 3, Fisher’s exact test p-value < 1.39e-02) (Fig. 4B, Tables S6 and S7). Unfortunately, all of ATEI transcripts with these variants do have more than one SNP in the transcript, thus it is impossible, short of generating congenic mouse lines, to definitively establish the precise causal variant(s). Nonetheless, some interesting possibilities emerge. Five variants in ATEI transcripts altered uORFs; 3 of them added a new uORF, and 2 of them disrupted an existing uORF. The disrupted uORFs did not affect mRNA expression, but were associated with either up- or down-regulated translation. For example, *Lyrm5* lost one uORF in FVB, correlating with increased TE (Fig. 4C), which indicates that the BL6 uORF may typically repress translation at downstream main ORF. Likewise, *Itgad* has one uORF in BL6 and lost it in FVB, which made no significant change in mRNA level but was associated with a −3.9 log fold change in translation (Fig. 4D). In addition, the phastCons conservation score is as high as 0.99, suggesting this sequence, and thus this uORF, has a conserved function on gene translation. Similarly, five ATEI genes have variants altering PAs in 3’ UTR, 2 of them gained an extra PAs and 3 of them lost a PAs. Among genes that are not differentially expressed in AEI, losing PAs enhanced their TE while gaining PAs reduced TE. For example, *Lyrm5* increased TE after losing one PAs (although this gene also lost one uORF, so it is unclear which change led to the TE increase), whereas *Lsm14b* and *Fam163a* had reduced TE after gaining one PAs. Overall, this suggests that variants impacting these features contribute to the differences in translation efficiency between these common alleles.

**Figure 4:**
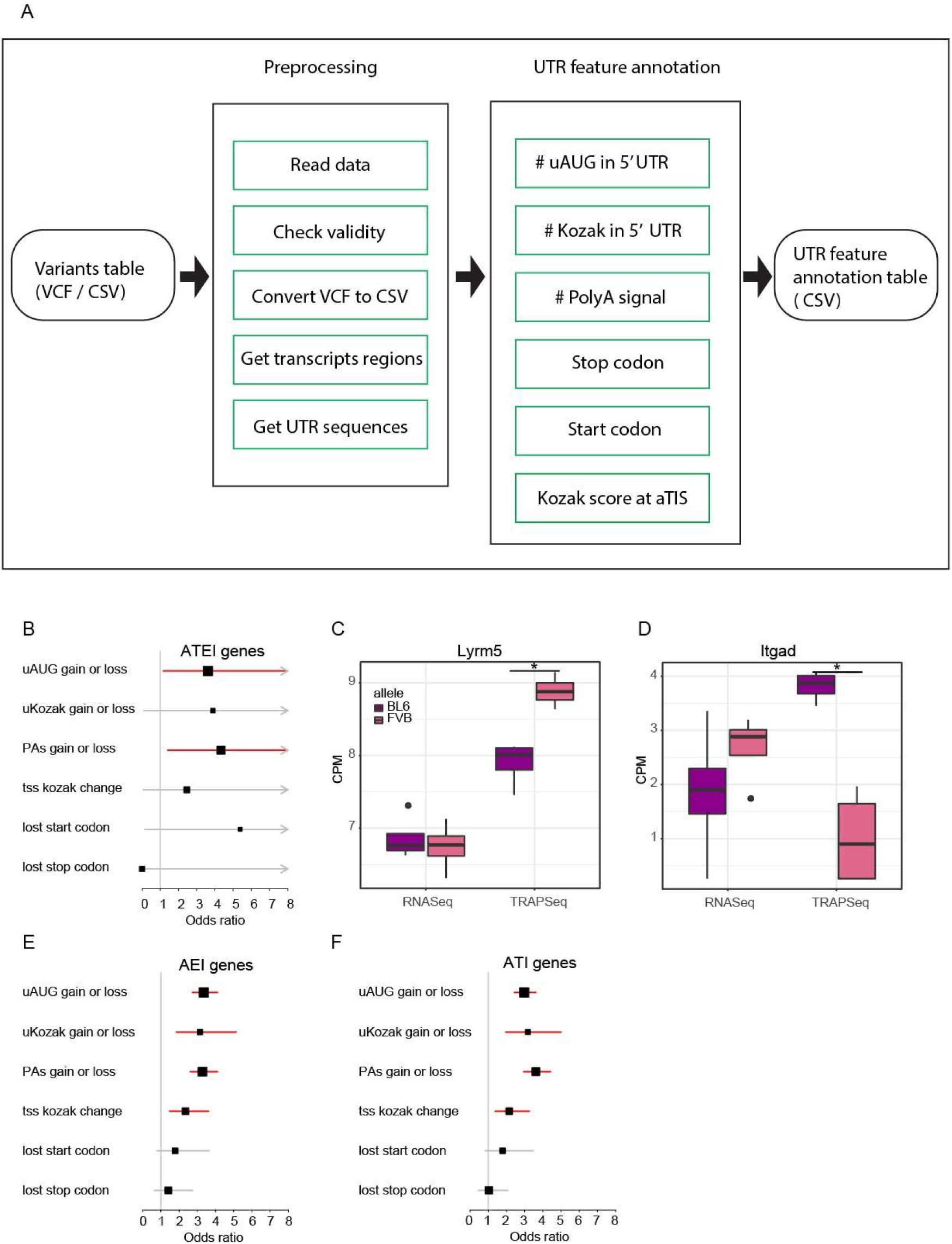
UTR variant annotator identifies significant enrichment of uORF and PAs altering-variants in AEI, ATI, and ATEI transcripts. A) Schematic of pipeline for UTR variant annotation (UTR pipeline 1.0). B) Forest plot showing odds ratios and confidence intervals testing whether variant classes are enriched in the ATEI transcripts. ATEI transcripts are enriched for variants that disrupt uORFs and PAs (*red lines*, p< .05). C,D) Examples of ATEI in the Lyrm5 and Itgad transcripts, which show a strong bias in TRAPSeq, but no or opposite changes in RNASeq (* p< 0.03). E,F) Forest plot for variant class enrichment in the AEI and ATI transcripts reveals 4 classes of variants (*red lines*, p< 6e-04) showing significant enrichment).

Though we had expected these kinds of variants would mostly influence ATEI genes, we were surprised to see a disproportionate amount of these events occurring in AEI (and the correlated ATI) transcripts as well. Variation that alters use of uORFs can influence stability, frequently by triggering nonsense-mediated decay (NMD) if a premature stop codon (PTC) is formed by their use (Mendell et al., 2004; Wittmann et al., 2006; Yepiskoposyan et al., 2011). Variation that alters polyA signals can cause truncations or extensions which introduce or remove elements that can alter a transcript’s post-transcriptional regulation, such as an AU-rich element, among others, and abnormally long 3’ UTRs that can arise from polyA variation are also triggers for NMD (Tian and Manley, 2017). Changes in mRNA stability can have a profound effect on the translational state of a transcript, and vice versa (Roy and Jacobson, 2013). Overall, we identify 126 AEI transcripts with uORF changes and 103 AEI transcripts with pAS changes. With each event having an Odds ratio of about 3, this suggests that 60 to 80 of the AEI differences are actually due to these SNPs impacting stability of the transcript as opposed to these SNPs(or others in linkage) altering transcriptional levels. Looking across these variants, however, we did not see a consistent direction of effect on RNA abundance for those variants altering predicted uORFs or PAs. Thus, these classes of variants appear to have equal chances of promoting or decreasing RNA stability overall.

### ATEI variants show higher impacts on predicted miRNA binding sites

Another key regulatory feature of UTRs is binding sites for miRNAs, so we also tested whether ATEI genes had increased SNPs falling in predicted miRNA binding sites. miRNAs are small (less than 35nt) RNAs that bind predominantly to 3’ UTR sequences, thereby recruiting Argonaute proteins to a transcript and leading to suppression of translation and potentially RNA decay (Rana, 2007). miRNA binding can be somewhat predicted by alignment, with key residues being bases 2-8 at the 5’ end. The miRanda prediction algorithm assigns scores to putative UTR target sequences based on a weighted average of all residues (Enright et al., 2003). We adapted this algorithm to calculate alignments for both alleles at all positions of variation in 3’ UTR between the mouse strains, and screened for variants that strongly altered the predicted binding (Fig. 5A). We compared these changes to a set of randomly selected control genes with matched length UTRs to the ATEI genes. Our first findings were that ATEI transcripts contain more SNPs than control transcripts (Fig. 5B), and these tend to fall in more conserved residues (Fig. 5C). Looking at the maximum absolute change of minimum free energy (MFE) of all transcripts, we found that ATEI transcripts had a significantly higher MFE change between alleles than control transcripts (Fig. 5D). This indicates that variants in ATEI UTRs tend to more strongly alter predicted miRNA::mRNA interaction. In many cases, some SNPs completely disrupted an miRNA binding site or introduced one (Fig. 5E, Fig 5F, Table S8). Finally, we repeated the analysis using miRNAs that were definitely expressed in astrocytes, leveraging our prior *in vivo* biochemical studies (Hoye et al., 2017; Liu and Wang, 2019). The same effect was true when limiting our analyses to these miRNAs (Fig. 5G). Thus, a subset of ATEI changes appear likely to be mediated by alterations in miRNA binding sites.

**Figure 5:**
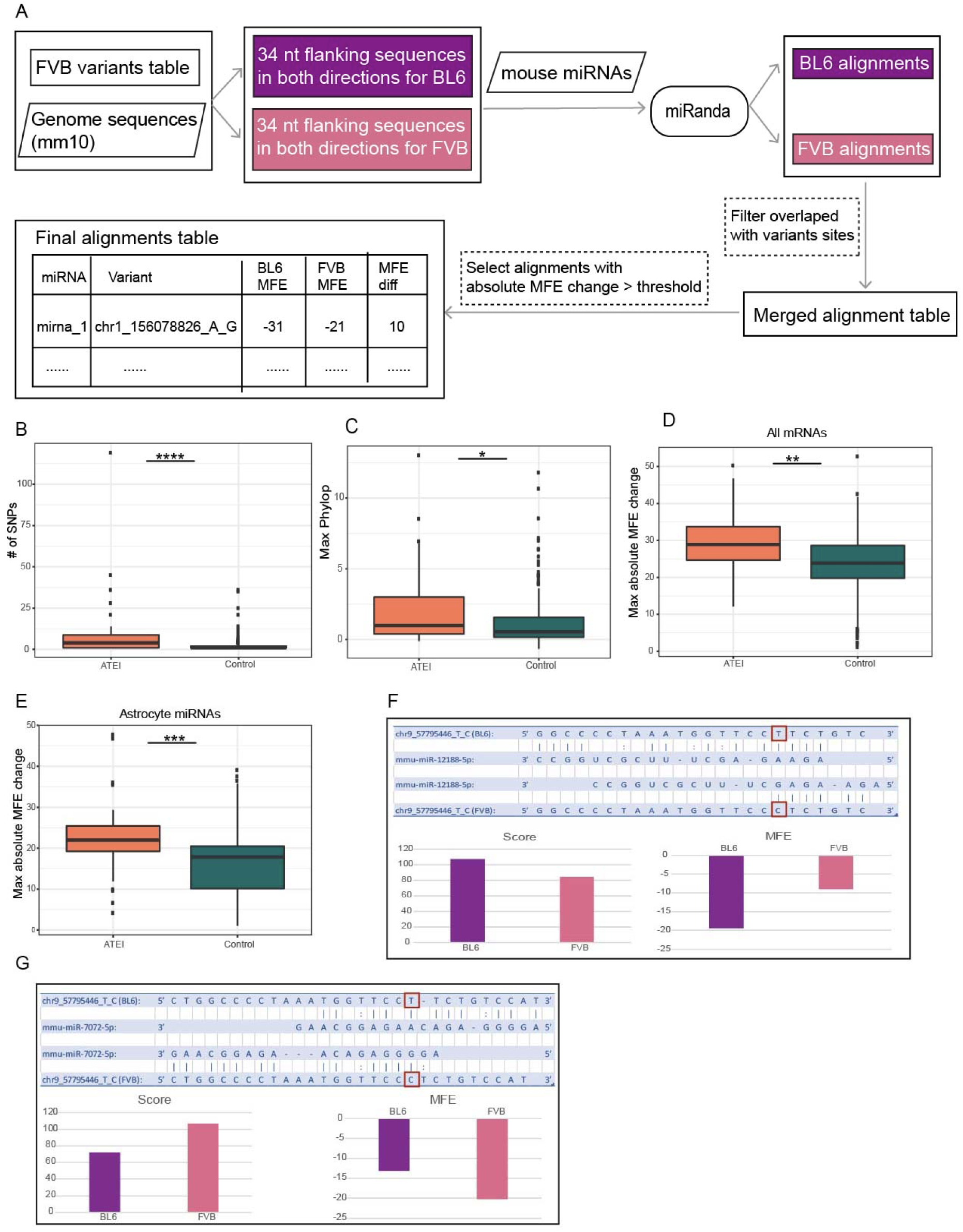
ATEI transcripts are associated with more variation and larger changes in predicted miRNA binding. A) Illustration of miRNA analysis pipeline. SNPs and surrounding sequence had their binding predicted for by all known miRNAs using miRanda. SNPs changing alignment scores for miRNAs were retained for downstream analysis. B). ATEI genes had a larger number of SNPs on average than matched length controls (*Wilcoxon, p=8e-11)*. C) SNPs in ATEI genes were in more conserved positions, as assessed by phyloP, than in control genes (*Wilcoxon, p=0*.*0097)*. D) ATEI transcripts had larger magnitude changes in predicted binding, illustrated as the maximal change in predicted minimum free energy for binding of miRNA to miRNA (*Wilcoxon, p=4*.*3e-06)*. E) Findings were similar when restricting to only those miRNAs measurably expressed in astrocytes (*Wilcoxon, p=1*.*3e-07)*. F,G) Example of SNPs altering predicted miRNA binding sites and corresponding free energy for Arid3b transcript.

## Discussion/Conclusions

SNPs and larger mutations can be deleterious to the proper regulation of transcript abundance and protein production, owing to the introduction or disruption of key elements, many of which lie outside the coding region for a given gene. In this work we sought to analyze how such effects could present in two closely related *Mus musculus* strains, as a proxy for how two humans who differ in genome sequence from one another may present differences in these processes. We established, in F1 progeny of a cross between C57BL/6J and FVB/NJ mice, that thousands of transcripts both differ in expression levels and differ in astrocyte-specific ribosome occupancy. While >90% of the ribosome occupancy as measured by TRAP could be explained by transcript levels, we found over a hundred ‘ATEI’ transcripts for which there was a significant difference between the change in expression vs. the change in ribosome occupancy, indicating a role for these variants beyond control of mRNA abundance. This suggests that common variants might alter translation efficiency in 1-2% of transcripts *in vivo*.

More detailed analysis of UTR elements revealed that within these ATEI genes, uORFs and poly-A signals were more likely to be disrupted than expected by chance. Additionally, many variants changed the MFE of miRNA binding sites, with several specifically potentially disrupting existing sites or introducing new sites. Also, those SNPs that were found in the ATEI transcripts were in positions that were more conserved when compared to those in control transcripts, suggesting these positions were indeed regulatory. Unfortunately though, only a minority of ATEI transcripts had just a single SNP in them, meaning in many cases it can be challenging to determine which variant in a linked block is causal. Regardless, many of the genes displaying ATEI also altered both AEI and ATI, indicating the translation efficiency is rarely regulated in a vacuum, and is consistent with prior observations indicating the translatability of a transcript can also alter transcript abundance.

It must be noted that while the ribosome-bound mRNA affinity purified from the brain homogenates was astrocyte specific (GFP-RPL10A expression being driven by the astroctye-specific *Aldh1l1* promoter) and used for ATI analysis, total brain RNA was used for AEI analysis and subsequent ATEI analysis. Moving forward, the parallel AEI studies could also be compartmentalized solely to a single cell type by first sorting brain homogenates or nuclei from TRAP lines by presence of GFP signal through fluorescence-assisted cell sorting (FACS) (Reddy et al., 2017). Such a sorting may remove any analysis artifacts that could arise through cell type-specific differences in expression, as there is significant variation in the transcriptome of different cell types (Delile et al., 2019; Jovičić et al., 2013; Shang et al., 2018; Zhong et al., 2018) and miRNA landscape (Jovičić et al., 2013), maintenance of which is critical for neural development. Likewise, it is interesting that the overall correlation between ribosome occupancy (measured by TRAP) and RNASeq was quite high (Figure 3D), and higher than expected from prior studies comparing RNASeq to Ribosome Footprinting(RF) or Polysome profiling, where the correlation has been substantially lower (Dalal et al., 2017; Ingolia et al., 2009, 2011). This suggests that TRAP could be underestimating the regulation due to translation and the 1-2% of transcripts showing ATEI due to common variants here might be a lower end estimate. Indeed, in mouse fibroblasts, Hou et al identified 14% of transcripts as showing ATEI, though that was comparing much more distally related mouse strains (∼35M SNPs diverged as opposed to 6M here), so it cannot be determined from this experiment alone. Head-to-head comparisons with the same cells and strains might be needed to see if there is indeed a difference in sensitivity between using TRAP compared to polysome profiling or RF. Regardless, it is clear that even if TRAP ends up less sensitive than polysome profiling for picking up ATEI in future benchmark studies, it is sufficiently sensitive to detect some changes and would continue to fill a unique niche with its ability to assess specific cell types *in vivo*.

By the same token, mRNA abundance analysis gives an idea of how variants alter a transcript’s expression, but it is unclear which process is being hindered or enhanced. Several methods for exploring nascent RNA production exist, such as capturing chromatin-associated RNA, Polymerase II-associated RNA, and metabolic labeling of newly synthesized RNA (Wissink et al., 2019). Additionally, measurement of RNA decay can be accomplished by either metabolic labeling of RNA and subsequent washout and time point monitoring of decay, or halting all transcription, such as with the use of actinomycin D, and collecting the time point samples. Combining these might allow a more comprehensive view of allelic effects at each step of an mRNAs life.

We do believe that the analytical approach presented here, of each sample into 2 ‘pseudo-samples’ by ancestry and using standard count-based packages for RNASeq analysis, presents an elegant solution to standardizing analysis for AEI, ATI, and ATEI in model organisms. This simplified approach should enable more labs to conduct screens to identify genetic regulators of translation, whether using TRAP, polysome profiling, or RF, and we have shared our pipeline as a repository on bitbucket as example code to facilitate future studies. Likewise, we hope that our pipeline for annotating UTR variants might also be of use for those interested in the potential consequences of mutations in UTRs, and we have likewise made this available for download.

Regardless of the technical considerations, we have shown here that non-coding SNPs can be associated with very strong changes in translational efficiency *in vivo*. At their most extreme levels, non-coding allelic variants that alter mRNA or final protein abundance sufficiently could give rise to effect levels similar to haploinsufficiency. Loss of a single copy of many highly constrained genes can result in severe disorders such as autism (Samocha et al., 2014). And UTR variants of sufficient effect might phenocopy such gene deletions. Outside of rare variants, most common SNPs statistically linked to disease that are discovered by genome-wide association studies (GWAS) map to non-coding regions (Zhang and Lupski, 2015). The non-coding region is being investigated especially in the context of neurodevelopmental disorders, and 3’ UTR variants have been implicated in autism spectrum disorder, intellectual disability, schizophrenia, and attention-deficit/hyperactivity disorder, among others (Wanke et al., 2018). Further investigation into allelic variants, especially *de novo* ones, and their impacts on expression regulation will help our understanding of their potential pathogenicity.

## Acknowledgements

We’d like to thank Krissy Sakers for images of astrocytes, and the Genome Technology Access Center (GTAC@MGI) in the Department of Genetics at Washington University School of Medicine for sequencing support. This work was supported by 5R01NS102272, 1R01MH116999, and the Simons Foundation (571009) to JDD, as well as P30 CA91842 (NCI) and UL1 TR000448 (NCATS) to GTAC@MGI.

## Figure and Table Legends

**Figure S1: Top Genes with ATEI**

Fifteen example genes displaying significant ATEI and having the lowest FDR corrected p-values. The RNASeq panels illustrate RNA expression imbalances, while the TRAPSeq panels depict translational (ribosomal occupancy) imbalances.

**Figure S2: Library Complexity Analysis**

VGAM package analysis of library complexity to assess bias. Dispersion values are indicated.

## Tables

**Table S1: Raw Gene Counts**

Raw gene counts for BL6 vs FVB variants.

**Table S2: Genes with AEI**

Genes with AEI having FDR corrected p-values <0.1. Gene ID from Ensembl, gene symbols, gene biotypes, Log2 fold change of CPM, Log2 CPM, likelihood ratio test scores, p-values, FDR corrected (Benjamini-Hochberg method) p-values, and number of SNPs in exons region. For Log2 FC, a positive number indicates read count was higher for the FVB variant, negative that read count was higher for the BL6 variant.

**Table S3: Genes with ATI**

Genes with ATI having FDR corrected p-values <0.1. Column headers and interpretation of FC are as in Table S2.

**Table S4: Genes with ATEI**

Genes with ATEI having FDR corrected p-values <0.1. Column headers and interpretation of FC are as in Table S2.

**Table S5: DE Analysis**

DE analysis results on AEI, ATI, and ATEI of all detectable genes. Gene ID from Ensembl, external gene names, gene biotypes, log2 FC, log2 CPM, likelihood ratio test scores, p-values, FDR corrected p-values of each analysis distincted by suffix, and number of SNPs in exons region.

**Table S6: ATEI Genes with Gained or Lost AUG Sequences**

ATEI genes which gained or lost AUG sequences, along with PhastCons and PhyloP scores, and log2 FC of CPM of ATEI, AEI, and ATI of the gene.

**Table S7: ATEI Genes with Gained or Lost PolyA Signals**

ATEI genes which gained or lost poly-adenylation signals, along with PhastCons and PhyloP scores, and logFC of CPM of ATEI, AEI, and ATI of the gene.

**Table S8: Top miRNA change by MFE for each ATEI transcript**

The top miRNA with maximum MFE changes between FVB and BL6 for each ATEI gene which only has 1 SNP in 3’ UTR. MFE_change is defined as the MFE of FVB - the MFE of BL6. MFE_change < 0 means the corresponding miRNA binds more stable in FVB than in BL6, and vice versa. Score_change is defined as the score of FVB - the score of BL6. Score_change < 0 indicates the corresponding miRNA aligns better in FVB than in BL6. Score, energy, and number of binding sites in FVB and BL6 are distinguished by the suffix (_FVB and _BL6). PhastCons and phyloP are average conservation scores of the mRNA binding regions.

