## Supplementary figures and images for "TRAP-based allelic translation efficiency imbalance analysis to identify genetic regulation of ribosome occupancy in specific cell types *in vivo*"

### Figure S1

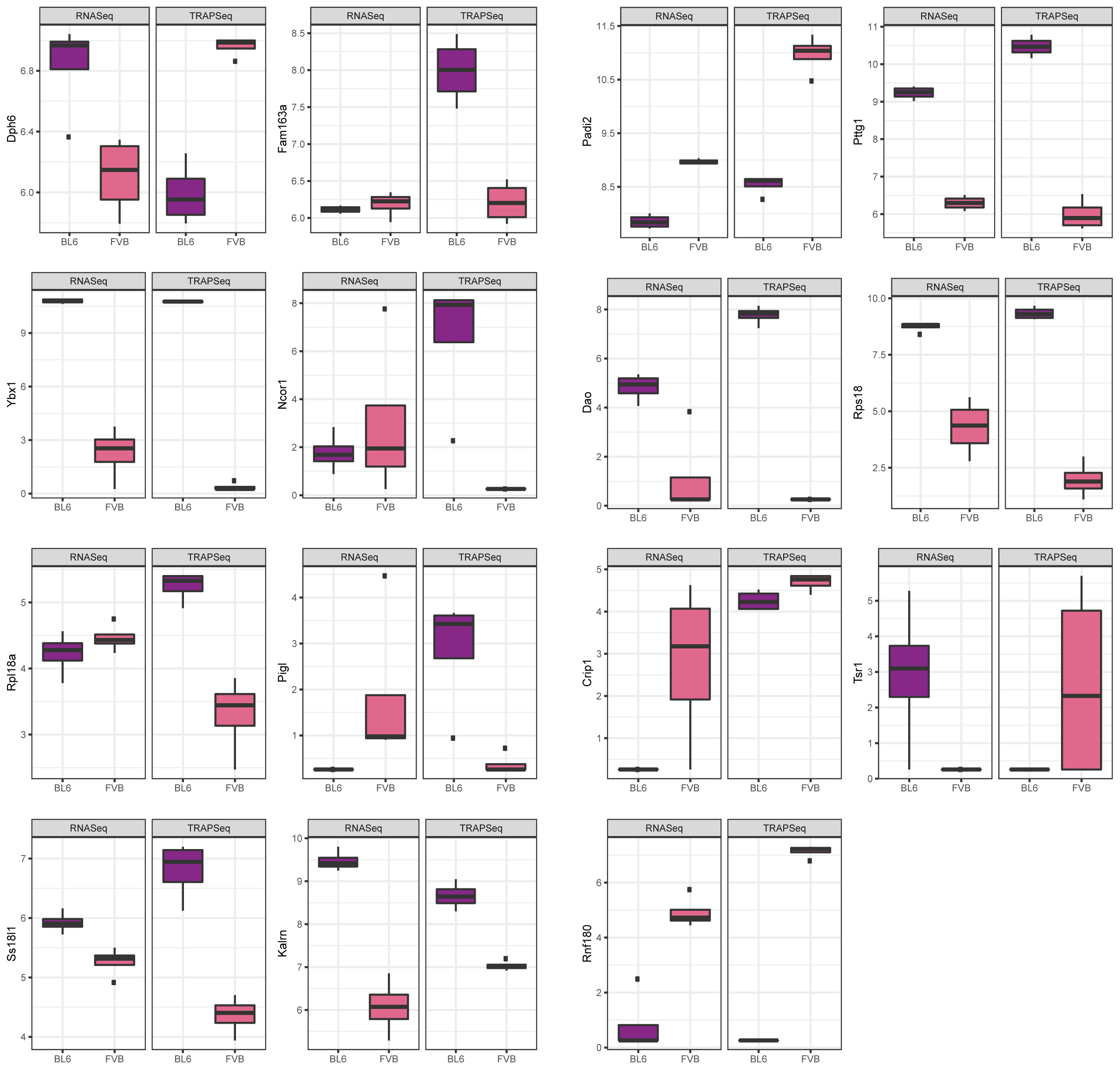

### Figure S2

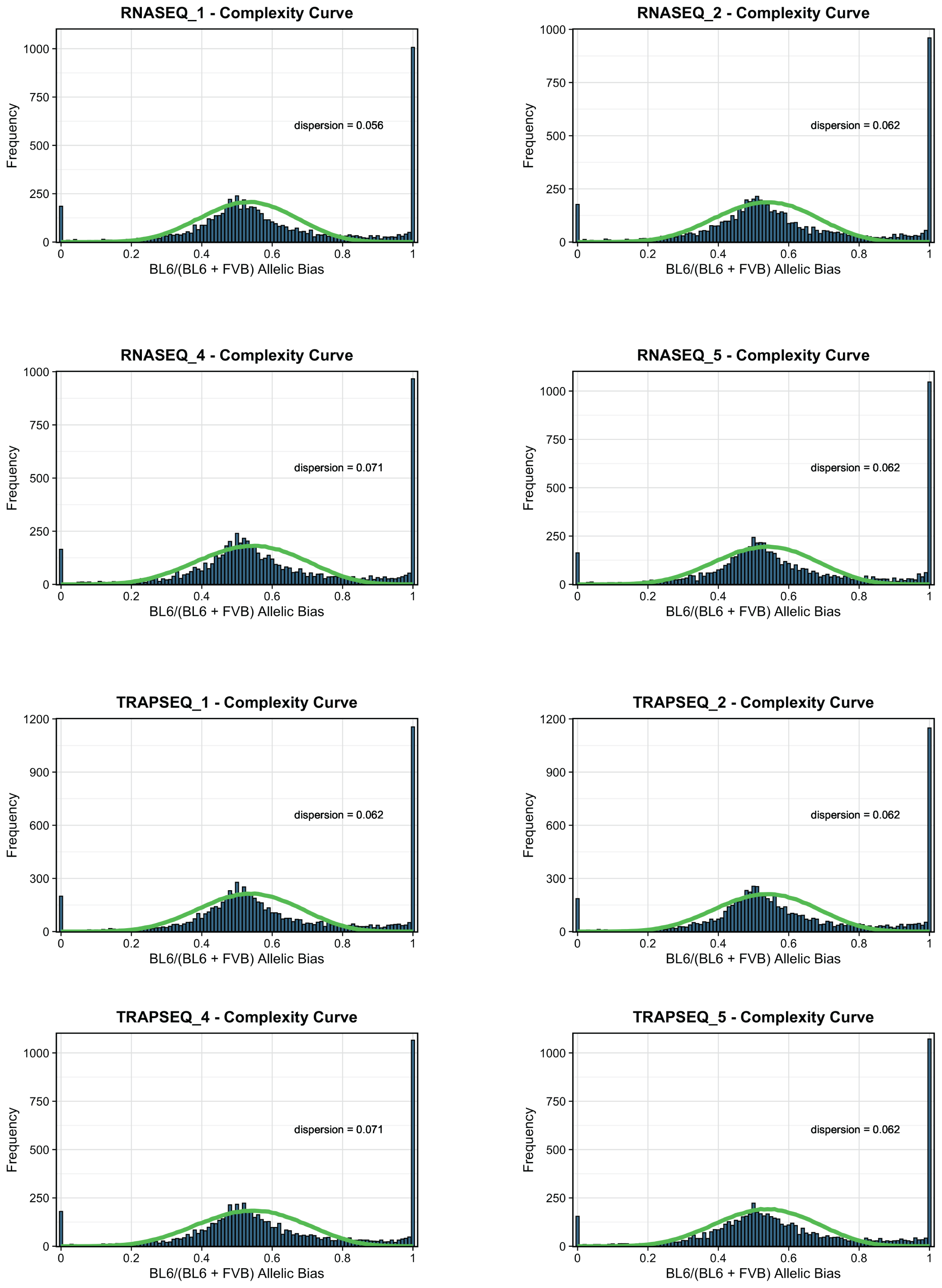
